# The odor of a nontoxic tetrodotoxin analog, 5,6,11-trideoxytetrodotoxin, is detected by specific olfactory sensory neurons of the green spotted puffers

**DOI:** 10.1101/2023.08.22.552781

**Authors:** Takehisa Suzuki, Ryota Nakahigashi, Masaatsu Adachi, Toshio Nishikawa, Hideki Abe

## Abstract

Toxic puffers accumulate tetrodotoxin (TTX), a well-known neurotoxin, by feeding on TTX-bearing organisms and using it to defend themselves from predators. Our previous studies have demonstrated that toxic puffers are attracted to 5,6,11-trideoxytetrodotoxin (TDT), a nontoxic TTX analog that is simultaneously accumulated with TTX in toxic puffers and their prey. In addition, activity labeling using immunohistochemistry targeting neuronal activity marker suggests that TDT activates crypt olfactory sensory neurons (OSN) of the green spotted puffer. However, it remains to be determined whether individual crypt OSNs can physiologically respond to TDT. By employing electroporation to express GCaMP6s in OSNs, we successfully identified a distinct group of oval OSNs that exhibited a specific calcium response when exposed to TDT in green spotted puffers. These oval OSNs showed no response to amino acids (AAs), which serve as food odor cues for teleosts. Furthermore, oval morphology and surface positioning of TDT-sensitive OSNs in the olfactory epithelium closely resemble that of crypt OSNs. These findings further substantiate that TDT is specifically detected by crypt OSNs in green spotted puffer. The TDT odor may act as a chemoattractant for finding conspecific toxic puffers and for feeding TTX-bearing organisms for effective toxification.

## Introduction

Tetrodotoxin (TTX) is a well-known neurotoxin found in toxic puffers that blocks sodium channels and prevents the generation and conduction of nerve impulses, leading to paralysis and potentially fatal respiratory and cardiac failure. Toxic puffers accumulate TTX in their bodies by feeding on TTX-bearing organisms and using it to defend themselves from predators (Noguchi et al. 2006). While some studies have suggested that TTX attracts toxic puffers through olfaction (Matsumura 1995; Okita et al. 2013), our electrophysiological and behavioral study has demonstrated that the grass puffer *Takifugu alboplumbeus* is attracted to 5,6,11-trideoxyTTX (TDT), a nontoxic TTX analog that is accumulated simultaneously with TTX in toxic puffer or their prey (Noguchi et al. 2022). Furthermore, our behavioral and immunohistochemical study also revealed that TDT attracts the green spotted puffer *Dichotomyctere nigroviridis*, which is phylogenetically and geographically quite different from the genus *Takifugu*, and then activates crypt olfactory sensory neurons (OSNs) (Suzuki et al. 2022). Based on these studies, we hypothesized that TDT, an attractive odorant in toxic puffers, is detected by crypt OSNs. However, we have yet to establish whether single crypt OSNs can physiologically respond to TDT.

In this study, we demonstrated the specific response of OSNs to TDT in green spotted puffers by expressing GCaMP6s, a genetically encoded calcium indicator. To achieve this, we used *in vivo* electroporation to perform transient exogenous gene transfer into the olfactory epithelium (OE), leveraging the unique anatomical features of the nasal organ of the green spotted puffer that protrudes from the body surface. The resulting TDT-sensitive GCaMP6s-expressing OSNs were oval cells located in the superficial layer of the olfactory epithelium, similar to the crypt-type OSNs previously identified by immunohistochemistry using neuronal activity marker antibody (Suzuki et al., 2022). These OSNs exhibited no response to amino acids (AAs) or TTX. Conversely, the AAs-sensitive OSNs were elongated or spindle-shaped cells, with their cell bodies positioned deeper within the olfactory epithelium than the TDT-sensitive OSNs. These AAs-sensitive OSNs did not respond to TDT, indicating a distinct detection mechanism in different OSNs. Various odorants are reported to act as ligands for crypt OSNs (Bazáes and Schmachtenberg 2012; Biechl et al. 2016; Schmachtenberg 2006; Suzuki et al. 2022; Vielma et al. 2008). Our findings provide direct evidence of TDT being specifically detected by crypt-like OSNs. Given our findings and that toxic puffers preferentially prey on TTX-bearing organisms (Ito et al. 2023; Itoi et al. 2015; Okabe et al. 2019), we speculated that TDT acts as a chemoattractant. TDT odor may aid toxic puffers in locating conspecifics and feeding on TTX-bearing organisms, thereby facilitating efficient toxification.

## Materials and Methods

### Animals

Green spotted puffers (*Dichotomyctere nigroviridis*; standard body length:1.5–3.0 cm, bodyweight:0.5–1.5 g) were purchased from a local dealer and maintained in 60 L tanks (28°C; 20–30 fish per tank) filled with artificial brackish water (ABW; Instant Ocean Sea-Salt, Cl^−^ 543 mM, Na^+^ 471 mM, Mg^2+^ 54.3 mM, K^+^ 10.7 mM, Ca^2+^ 10 mM, Br^−^ 0.7 mM, Sr^2+^ 0.1 mM, F^−^ 0.05 mM, I^−^ 0.002 mM, CO_3_^2−^/HCO_3_^−^ 3.3 mM, pH 8.2; Instant Ocean, Blacksburg, VA) with a specific gravity of 1.010 under a 12 h light/12 h dark photoperiod. The fish were fed dried shrimp once a day. Since it is difficult to determine sex and sexual maturity from appearance, several fish were anesthetized with tricaine methane sulfonate (MS-222; 0.02%, Sigma-Aldrich, St. Louis, MO), and their gonads were dissected. All fish identified were juveniles with immature gonads; therefore, both males and females were used without distinction in this study. The animal use and care protocols were approved by the Center for Animal Research and Education of the Nagoya University (approved number: A210871-004). All animals were maintained and used in accordance with the Tokai National Higher Education and Research System Regulations on Animal Care and Use in Research (Regulation no. 74). No statistical methods were used to predetermine the sample size because many of the outcomes were unknown.

### Plasmid

The plasmid containing the medaka β-actin promoter and GCaMP6s was constructed by modifying of *pT2Olactb:loxP-dsR2-loxP-EGFP* plasmid (Yoshinari et al. 2012; a gift from Dr. Masato Kinoshita). The dsR2-loxP-EGFP sequence between the Spe I and Not I sites of *pT2Olactb:loxP-dsR2-loxP-EGFP* was replaced with the *GCaMP6s* sequence, which was obtained as Spe I and Not I fragments from *pGP-CMV-GCaMP6s*. *pGP-CMV-GCaMP6s* was a gift from Douglas Kim & GENIE Project (Chen et al. 2013; Addgene plasmid # 40753; http://n2t.net/addgene:40753; RRID:Addgene_40753). The purified plasmid was prepared using the NucleoBond® Xtra Midi kit (MACHEREY-NAGEL, Düren, Germany) or the Genopure Plasmid Midi kit (Roche Diagnostics GmbH, Mannheim, Germany). The prepared plasmid was dissolved in TE buffer to 5 µg/µL and stored at −20°C until use.

### *In vivo* electroporation to the olfactory epithelium of green spotted puffer

The fish were first anesthetized with 0.02% MS-222, and excess water was wiped off. Then, 2 µL of a plasmid solution (1 µg/µL) diluted in PBS was administered. The olfactory organ was positioned between self-made tweezer-style silver chloride/silver plate electrodes. The gap between the electrodes was adjusted to 2-3 mm, and square-current pulses were delivered to the olfactory organ using an electronic stimulator (SEN-7203; Nihon Kohden, Tokyo, Japan) through an isolator (SS-202J; Nihon Kohden) at room temperature.

A two-step pulse protocol was used for gene transfer. This protocol comprised an initial high-current single-shot poring pulse with parameters of amplitude (10 mA) and duration (5 ms), followed by multiple low-current transfer pulses (**Fig. 1A**). The pulse parameters, including the interval between the poring and transfer pulse patterns (20 ∼ 1000 ms), amplitude (0.5 ∼ 2.0 mA), and frequency (5 ∼ 20 Hz) of the transfer pulses, were optimized as described below.

**Fig. 1:**
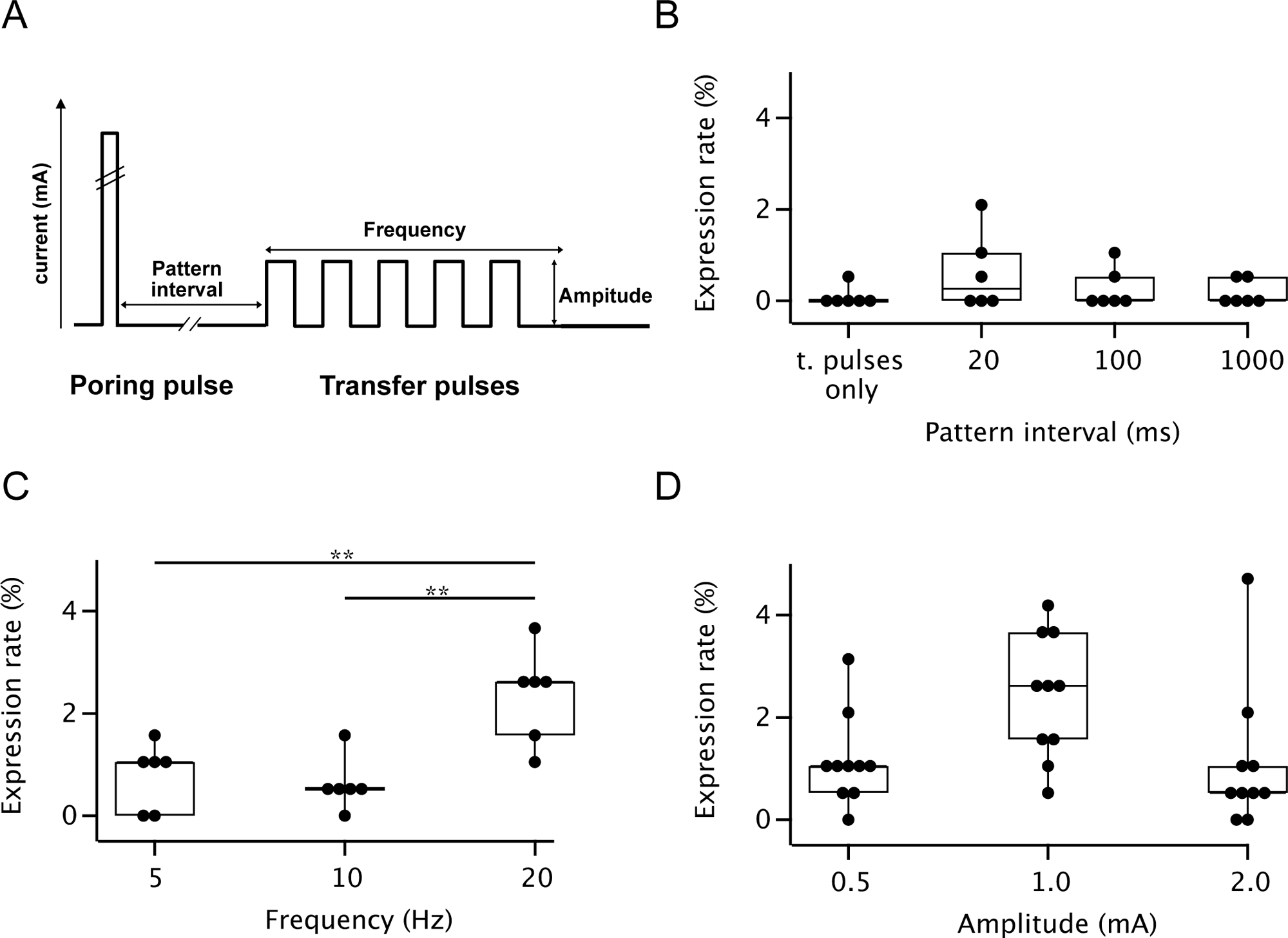
*In vivo* electroporation of GCaMP6s to OSNs. Schematic diagram representing the waveform used for electroporation (**A**). Effects of the pattern interval between the poring and the transfer pulse (**B**), transfer pulse frequency (**C**), and transfer pulse amplitude (**D**) on the expression rate of GCaMP6s in OSNs. Each condition was repeated for six trials (for pattern interval and transfer pulse frequency) or ten trials (for transfer pulse amplitude). Dunnett’s multiple comparison test (in comparison with t. pulses only; for (**B**)) or Tukey’s multiple comparison test. **P < 0.01.

Following electroporation, the fish were allowed to recover for 24-36 h in fresh ABW before use in the experiments. Gene transfer performance was evaluated as the rate of OSNs expressing GCaMP6s per OSNs counted using 4,6-diamidino-2-phenylindole (DAPI) staining normalized to the OE area (90×90 µm^2^; average OE cell density is 191 cells, n = 4). Cell counting was performed with 3D segmentation and the 3D ROI Manager function of the ImageJ plugin 3D suite (Ollion et al. 2013). To ensure accurate quantification of living OSNs, we included only those cells located within the sensory epithelial layer and exhibiting GCaMP6s fluorescence. Cells that overexpressed fluorescent proteins or exhibited any morphological changes indicative of damage or cell death were excluded.

### Odorant solutions

A mixture of amino acids (AAs), including L-arginine, L-glutamine, L-phenylalanine, and L-isoleucine (Sigma-Aldrich), was dissolved in ultrapure water to a concentration of 10^−4^ M and stored as a stock solution at −30°C until use. TDT was synthesized and purified by HPLC using an ion-exchange column (Hitachi gel 3013C). The synthesized TDT was confirmed free of other TTX analogs and reagents, as verified by their ^1^H and ^13^C NMR spectra (**Fig. S1**). The purification procedure has been outlined in detail by Adachi et al. (2014). TDT and TTX (for biochemistry, citrate buffered, FUJIFILM Wako Pure Chemical Corporation) were dissolved in ultrapure water at 10^−3^ M concentration and stored as a stock solution at −30°C until use.

For puff application, all stock solutions were diluted with artificial cerebrospinal fluid for teleosts (ACSF) comprising NaCl (140 mM), KCl (5 mM), MgCl_2_ (1.3 mM), CaCl_2_ (2.4 mM), HEPES (10 mM), and glucose (10 mM), and adjusted to pH 7.4 to obtain a working concentration of 10^−6^ M.

### Calcium imaging

Electroporated fish were anesthetized with 0.02% MS-222, and the bilateral olfactory organs were excised and halved with a fine blade. The halved olfactory organ was transferred to a recording chamber and anchored with a handmade platinum slice anchor with parallel nylon threads (0.040 - 0.049 mm thickness; C-23S-N5 #8-0, Natsume-Seisakusyo, Tokyo, Japan). The preparation was continuously perfused with ACSF at a rate of 1 mL/min using a peristaltic pump (SJ1211-L, ATTO, Tokyo, Japan) and a vacuum pump (DAL-5s, ULVAC KIKO, Inc., Miyazaki, Japan).

A puff micropipette (with a tip diameter of 2 µm) filled with an odorant solution was carefully positioned 30-50 µm away from the OSN expressing GCaMP6s using a micromanipulator (EMM-3SV, Narishige, Tokyo, Japan) under a fluorescence microscope (Eclipse E600-FN, Nikon, Tokyo, Japan). The microscope was equipped with a 60× water immersion objective lens (CFI Fluor 60X/1.00W; Nikon), a fluorescence filter cube (excitation filter 450∼490nm, dichroic mirror 505 nm, and emission filter 520 nm; model B-2A, Nikon), and a CMOS camera (Chamelon3 CM3-U3-31S4M-CS, Teledyne FLIR LLC., Wilsonville, OR).

Puff application of the odorant solutions to OSN was performed by applying air pressure (0.8 kg/cm^2^; 500 ms duration; generated by an air compressor) to the puff micropipette using a solenoid valve (UMG1-T1, CKD, Komaki, Aichi, Japan). Time-lapse image acquisition (2 frames/s for 90 s) and puff application were controlled by Micromanager (Edelstein et al. 2014) and Arduino UNO (http://Arduino.cc). The morphologies of GCaMP6s-expressing OSNs for calcium imaging were confirmed through epifluorescence microscopy before starting the recording session. In some specimens that retained GCaMP6s fluorescence with minimal photobleaching after the experiment, detailed morphology was further confirmed by acquiring fluorescence optical sections (Z-stack) using a confocal laser microscope (FV1000D IX81, Olympus, Tokyo, Japan). The fluorescence change was calculated as ΔF/ F_0_= (F_t_ – F_0_)/F_0_, where F_t_ is the fluorescence intensity at a given time point, and F_0_ is the baseline (the average intensity of the first 10 frames). The criteria for determining a response in an OSN to the odorant puff stimulation was established as follows: if the OSN exhibited an increase in GCaMP6s fluorescence intensity, quantified as a change in ΔF/F0, of more than 3% within a 40-second timeframe following the puff stimulation of the odorant (AAs or TDT). Time-lapse images were analyzed using Fiji (https://fiji.sc/; Schindelin et al. 2012) and Igor Pro 9.01 (WaveMetrics, Portland, OR).

### Image analysis

Fluorescent optical sections of the non-fixed OE electroporated with GCaMP6s gene were acquired using an FV1000D IX81 confocal laser-scanning microscope (Olympus). Acquired Z-stack images of GCaMP6s fluorescence were preprocessed using Fiji to reduce confocal noise, employing a median filter with a radius of 2. Three-dimensional reconstruction and detailed visualization of the morphology of GCaMP6s-expressed OSNs were performed using FluoRender (http://www.sci. utah.edu/software/fluorender.html; Wan et al. 2017). FluoRender, an open-source interactive rendering tool, is designed to facilitate the visualization of confocal microscopy data. Adjustments to brightness, contrast, and gamma settings were also made using FluoRender to ensure the morphology of individual OSNs was visually discernible (**Fig. 2C, 3C**, and **3F**). The final arrangement of figures was completed using Affinity Designer ver.1.10.8 (Serif Europe Ltd., West Bridgford, Nottinghamshire, UK).

**Fig. 2:**
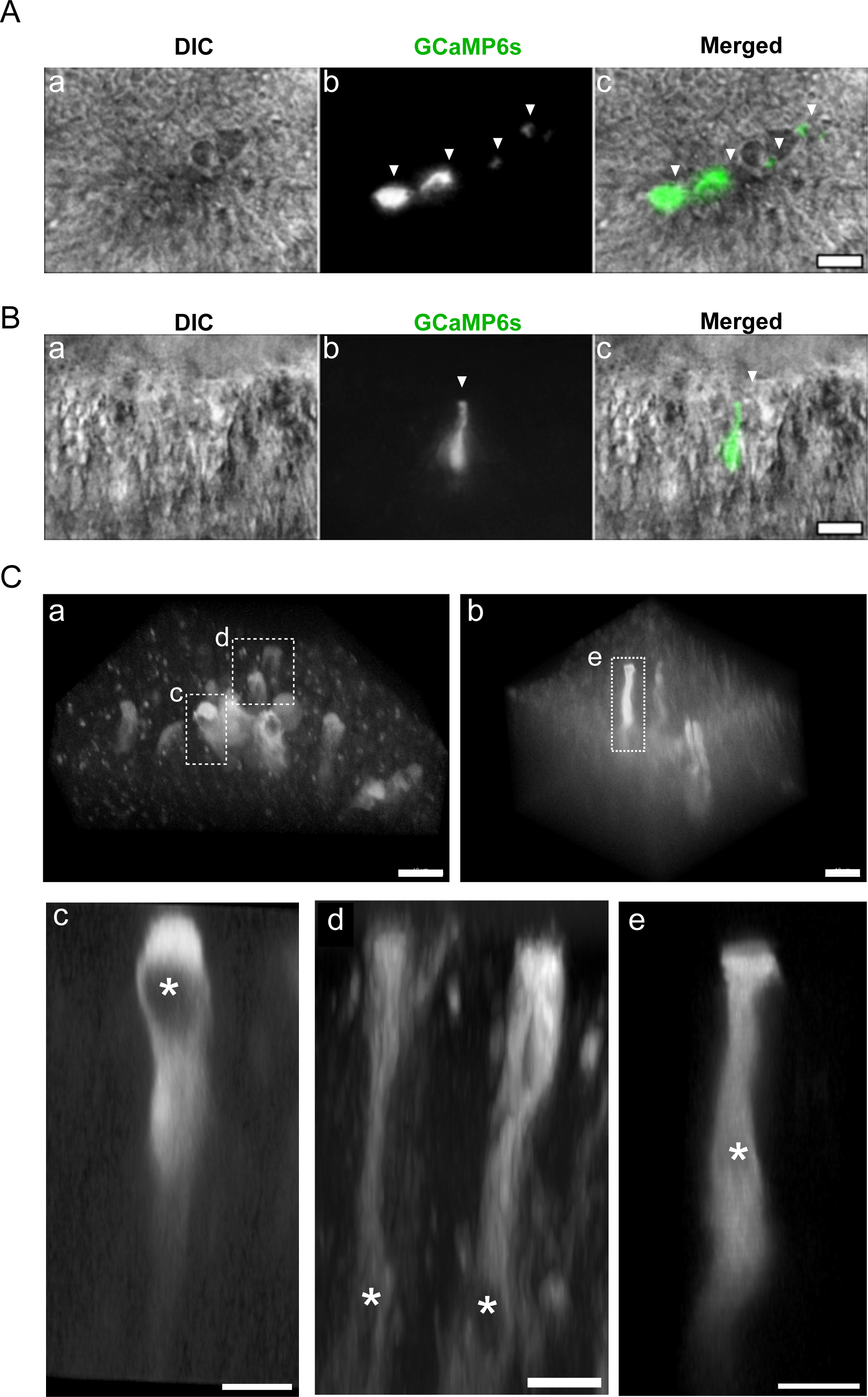
GCaMP6s-expressing OSNs in green spotted puffer’s OE following electroporation using our employed protocol. (**A**) Distribution of the GCaMP6s-expressing cells in the electroporated OE. Differential interference contrast (DIC; **a**), epifluorescence (**b**), and a merged image of the OE surface of the OE surface from the green spotted puffer (gray: DIC, green: GCaMP6s, **c**). The GCaMP6s-expressing cells are dispersed in the OE, allowing for the visualization of individual OSN’s morphology. Arrowheads indicate GCaMP6s expressing OSNs. (**B**) Pseudo-cross-sectional view of another electroporated OE. DIC (**a**), epifluorescence (**b**), and a merged image of the OE (gray: DIC, green: GCaMP6s, **c**). In this image, a GCaMP6s-expressing OSN extend an apical dendrite from the cell body in the middle layer of the OE to the surface. Arrowheads indicate GCaMP6s-expressing OSN. (**C**) Confocal laser-scanning microscopy images of the GCaMP6s-expressed OSNs. Bird’s-eye view volume-rendering images of the electroporated OE surface created from Z-stacks of confocal laser scanning microscopy. Two images from different OEs are shown in (**a**) and (**b**). The dashed rectangles in the images indicate the OSNs positions shown in **c∼e**. Volume-rendering x-z images of a single GCaMP6s-expressing OSN generated from (**a**) or (**b**) are shown in (**c∼e**): An oval OSN without apical dendrites in the surface layer of OE (**c**); a spindle-shaped OSN extending a swollen apical dendrite from the cell body in the middle layer of OE (**d**); an elongated OSN extending a thick and short apical dendrite from the cell body in the middle layer of OE (e). Asterisks in c∼e indicate the location of the nucleus. Scale bars: 10 µm (**A**, **B**, **Ca**, and **Cb**) and 5 µm (**Cc** to **Ce**).

**Fig. 3:**
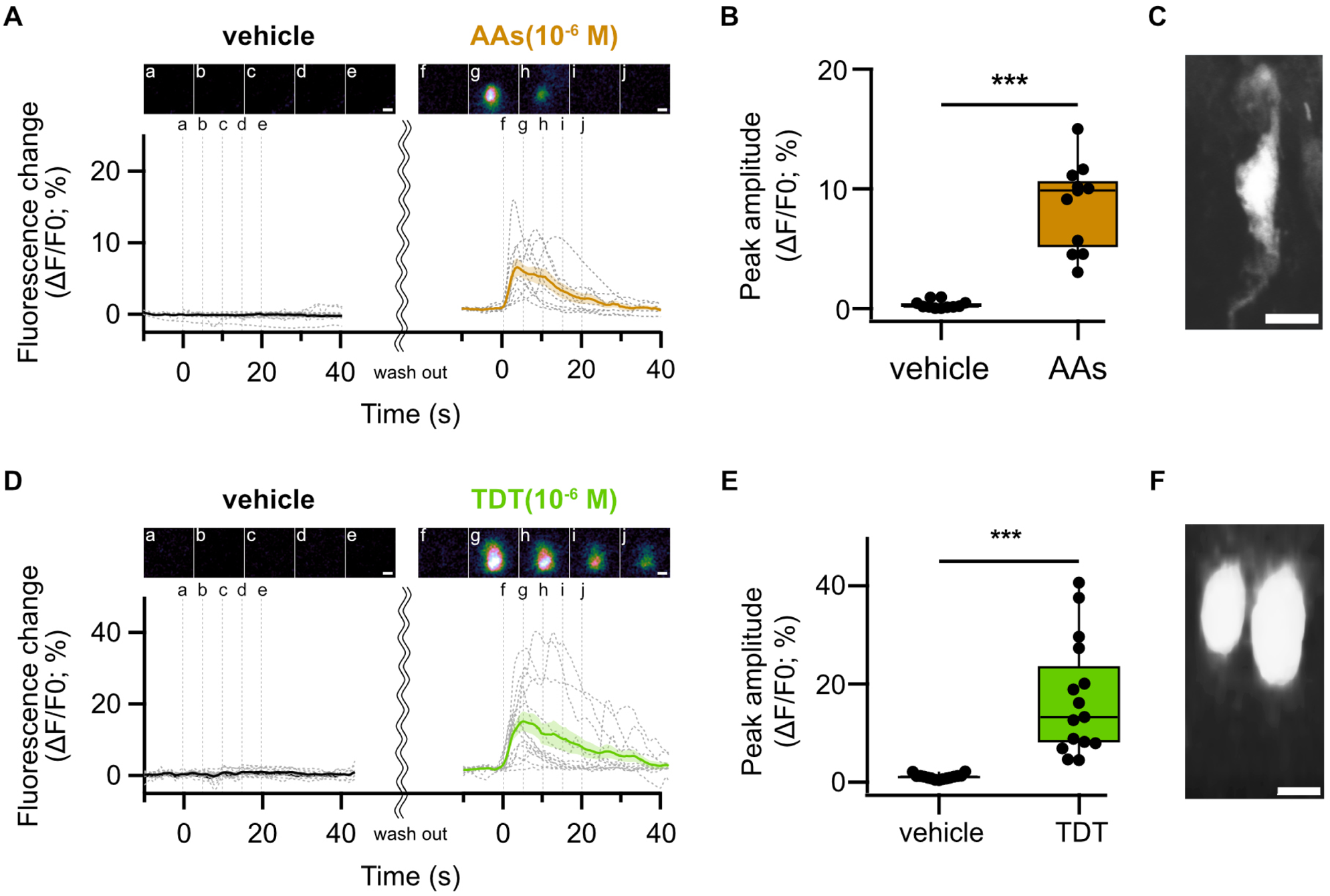
Response of OSNs to AAs or TDT. (**A**) Fluorescence changes in AAs-sensitive OSNs to the vehicle or AAs exposure. Individual AAs-sensitive OSNs (n = 14) are represented by gray dashed lines, with the bold line and envelope showing the mean ± SEM. Representative fluorescence image frames (**a-e**; the vehicle, **f-j**; AAs) indicated by vertical lines are shown above the trace. (**B**) The peak amplitude of AAs-sensitive OSNs after administering the vehicle (black) or AAs (brown). Paired t-test compared the vehicle vs. AAs administration. ***P < 0.001. (**C**) Volume-rendered x-z image generated from z-stacks of the confocal laser-scanning microscopy of the AAs-sensitive OSN as depicted in **A**. Scale bar: 5 µm. (**D**) Fluorescence changes in TDT-sensitive OSNs to the vehicle or TDT exposure. Individual TDT-sensitive OSNs (n = 15) are represented by gray dashed lines, with the bold line and envelope showing the mean ± SEM. Representative frames (**a-e**; the vehicle, **f-j**; TDT) indicated by vertical lines are shown above the trace. (**E**) The peak amplitude of TDT-sensitive OSNs after administering the vehicle (black) or TDT (green). Paired t-test compared the vehicle vs. TDT administration. ***P < 0.001. (**F**) Volume-rendered x-z image generated from z-stacks of the confocal laser-scanning microscopy of the TDT-sensitive OSN as depicted in **D**. Scale bar: 5 µm.

### Statistical Analysis

Data are expressed as the mean ± standard error of the mean and represented as a bee swarm plot and box-whisker plot (marker: mean; box: median and quartiles; whiskers: 10%–90% range) or a line plot. Igor pro 9 (WaveMetrics Inc., Lake Oswego, OR) and JASP (JASP team, 2023; Version 0.17.2) were used for statistical analysis and graph preparation.

## Results

### Optimization of Electroporation Conditions for Transient Gene Transfer to Olfactory Sensory Neurons

To investigate the physiological response of OSNs to TDT, we used a transient gene transfer method using *in vivo* electroporation to express GCaMP6s in the OSNs of green spotted puffers. Initially, we applied a train of square-current pulses (5 pulses, 50 ms duration, 10 Hz frequency, and 0.5 mA current amplitude; transfer pulse; **Table 1**) to the OE of green spotted puffers following dropwise addition of the GCaMP6s plasmid solution. The resulting GCaMP6s expression rate (the percentage of the number of GCaMP6s-expressing cells in the total olfactory epithelial cells of a 90 µm x 90 µm area) was 0.09 ± 0.01% (n = 6; measured 24∼36h after electroporation; **Fig. 1B**).

**Table 1:**
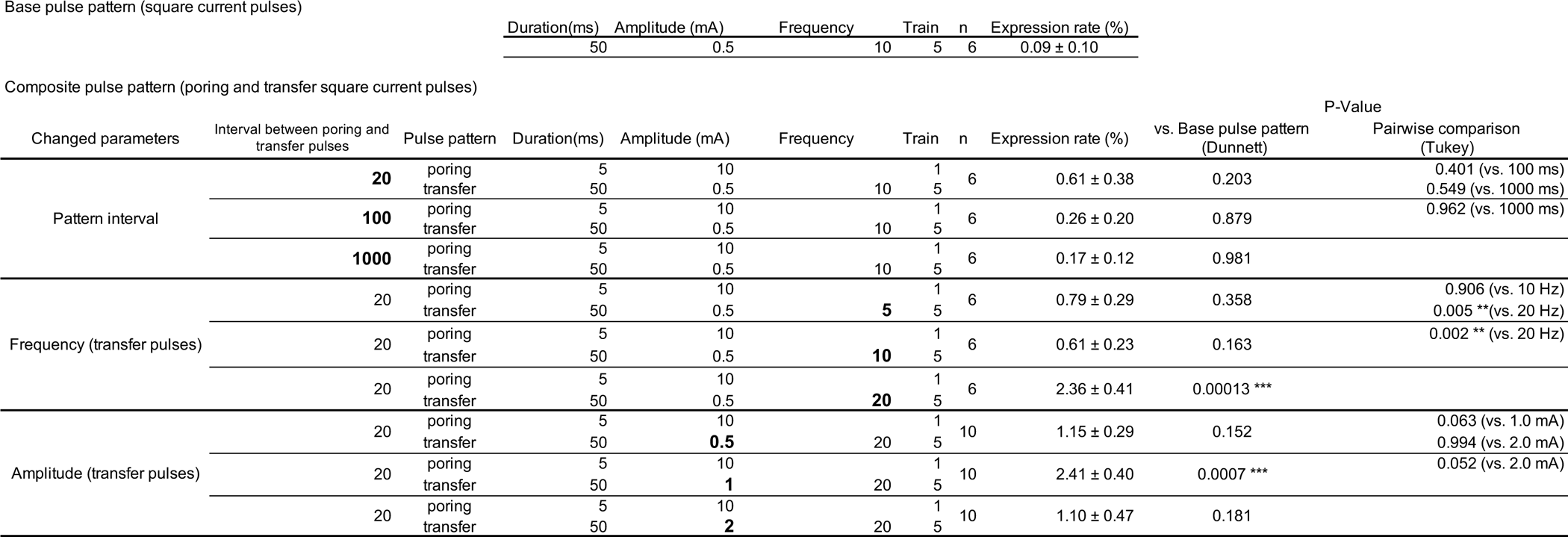
Electroporation pulse condition and GCaMP6s expression rate. Bold indicates the modified parameters.

To enhance the efficiency of gene transfer, we implemented a two-step current-pulse protocol consisting of an initial high-current, single-shot poring pulse (10 mA, 5 ms) followed by five low-current transfer pulses (0.5 mA, 50 ms, and 50% duty cycle; **Fig. 1A**). By adjusting the interval between the two pulse patterns, we observed an increase in the GCaMP6s expression rate but no significant difference was observed (**Fig. 1B**). Therefore, we connected the poring pulse and transfer pulses at an interval of 20 ms and set the poring pulse waveform (10 mA, 5 ms) and transfer pulse duty cycle (50%) to be constant for subsequent experiments.

Next, we investigated the effect of the transfer pulse frequency by changing the pulse duration while maintaining the amplitude of the transfer pulse at 1.0 mA. The GCaMP6s expression rate at 20 Hz transfer pulse frequency was significantly higher than those of 5 Hz or 10 Hz (**Fig. 1C**). Additionally, when we varied the amplitude of the transfer pulse while maintaining the poring pulse waveform and the frequency of the transfer pulse constant, the GCaMP6s expression rate tended to increase at 1.0 mA, although the increases were not statistically significant. However, further increasing the transfer pulse amplitude to 2.0 mA led to a decrease in the expression rate and resulted in the rupture and death of many OSNs (**Fig. 1D**). The changes in GCaMP6s expression rate under these conditions are summarized in **Table 1**.

Based on these results, we employed a two-step pulse protocol consisting of a 5 ms poring pulse with a 10 mA current amplitude, followed 20 ms later by five transfer pulses with a 1.0 mA current amplitude and 25 ms duration at 50% duty cycles. *In vivo* electroporation using these parameters resulted in scattered GCaMP6s-expressing OSNs on the OE of green spotted puffers at a density where individual cell shapes could be distinguished under epifluorescence microscopy (24∼36 hours post-electroporation; arrowheads in **Fig. 2A and B**). Since one side of the petal-like nasal organ has a mortar-shaped concavity, pseudo-cross-sectional planes of the OE can be obtained by focusing on its surface. Observing such pseudo-cross-sectional planes enabled us to distinguish the shape of individual GCaMP6s-expressing OSNs and their positions within the OE before the Ca^2+^ imaging (**Fig. 2B and S2A**). Furthermore, by comparing the detailed morphologies using volume-rendered images generated from z-stacks of confocal laser-scanning microscopy (**Fig. 2C**), we broadly classified the observed GCaMP6s-expressing OSNs into two types: oval cells lacking apical dendritic structures, with the cell body located in the surface layer of the OE (**Fig. 2Cb**), and elongated or spindle-shaped cells featuring swollen apical dendrites, with the cell body located in the middle layer of the OE (**Fig. 2Cc and Cd**). The oval OSNs had apical invagination and the processes extending from those cells, and the nuclei of oval OSNs (the compartment missing cytoplasmic GCaMP6s fluorescence) were located in the lower side of the soma.

### OSNs exhibit specific calcium responses to amino acids or TDT

Using electroporated OE, we investigated the physiological response of individual OSNs to odorants by detecting fluorescence changes in GCaMP6s. Puff administration of a 10^−6^ M mixture of four amino acids (L-Arginine, L-Glutamine, L-phenylalanine, and L-Isoleucine; AAs), which are known to act as odorants in many fish, induced a transient intracellular Ca^2+^ increase in 20 of 104 GCaMP6s-expressing OSNs. Among them, response to ACSF used as a vehicle control was also recorded from 11 AAs-sensitive OSNs, but no response was observed (**Fig. 3A**). The peak amplitudes of the Ca^2+^ response (expressed as ΔF/F_0_; %) induced by AAs significantly greater than those of the vehicle (8.9 ± 1.1% for AAs vs. 0.3 ± 0.3% for the vehicle (n = 11), P = 0.000018; **Fig. 3B**), and the repetitive AAs administration induced consistent Ca^2+^ responses (**Fig. S3A**). The volume-rendered x-z images generated from z-stacks of confocal laser-scanning microscopy after the calcium imaging exhibited the AAs-responsive OSNs had spindle-shaped morphology with a short diameter of approximately 4 μm. Their cell bodies were located slightly deeper on the OE surface (**Fig. 3C**). Twenty-three of 148 TDT-sensitive OSNs also exhibited a transient increase in Ca^2+^ (**Fig. 3D**). Among them, vehicle responses could also be recorded from 15 OSNs, the amplitude of TDT responses were significantly higher than the vehicle’s (17.1 ± 3.0% for TDT vs. 1.17 ± 0.1% for the vehicle, n = 15, P = 0.00012; **Fig. 3 E**), and the repetitive TDT administration induced consistent Ca^2+^ responses (**Fig. S3B**). The volume-rendered x-z images generated from z-stacks of confocal laser-scanning microscopy exhibited that TDT-sensitive OSNs exhibited oval-shaped morphology with a short diameter of approximately 6 μm, and their cell bodies were located on the surface of the OE (**Fig. 3F**). Cells expressing GCaMP6s, which lack apical dendrites and are located on the OE surface, could also be identified before calcium imaging using epifluorescence microscopy (**Fig. S2A**). These cells consistently exhibited calcium responses to TDT administrations (**Fig. S2B**). TDT-sensitive OSNs exhibited no fluorescence increase following the TTX application (**Fig. S4**).

We further examined the response of TDT-sensitive OSNs to AAs and vice versa. Regardless of the order of application, TDT-sensitive OSNs did not exhibit any fluorescence increase following the application of the vehicle or AAs (0.9 ± 0.1% for the vehicle vs. 8.5 ± 2.1% for TDT, P = 0.005; the vehicle vs. 0.7 ± 0.2% for AA, P = 0.901; TDT vs. AAs, P = 0.005, n = 6, respectively; **Fig. 4A and B**). Similarly, AAs-sensitive OSNs did not display any fluorescence increase by application of TDT or the vehicle (0.3 ± 0.1% for the vehicle vs. 0.3 ± 0.1% for TDT, P = 0.786; the vehicle vs. 13.6 ± 1.3% for AAs, P = 0.00000056; TDT vs. AAs, P = 0.00000056, n = 6, respectively; **Fig. 4C, and D**). Notably, no OSNs were observed to respond to both odorants, AAs and TDT.

**Fig. 4:**
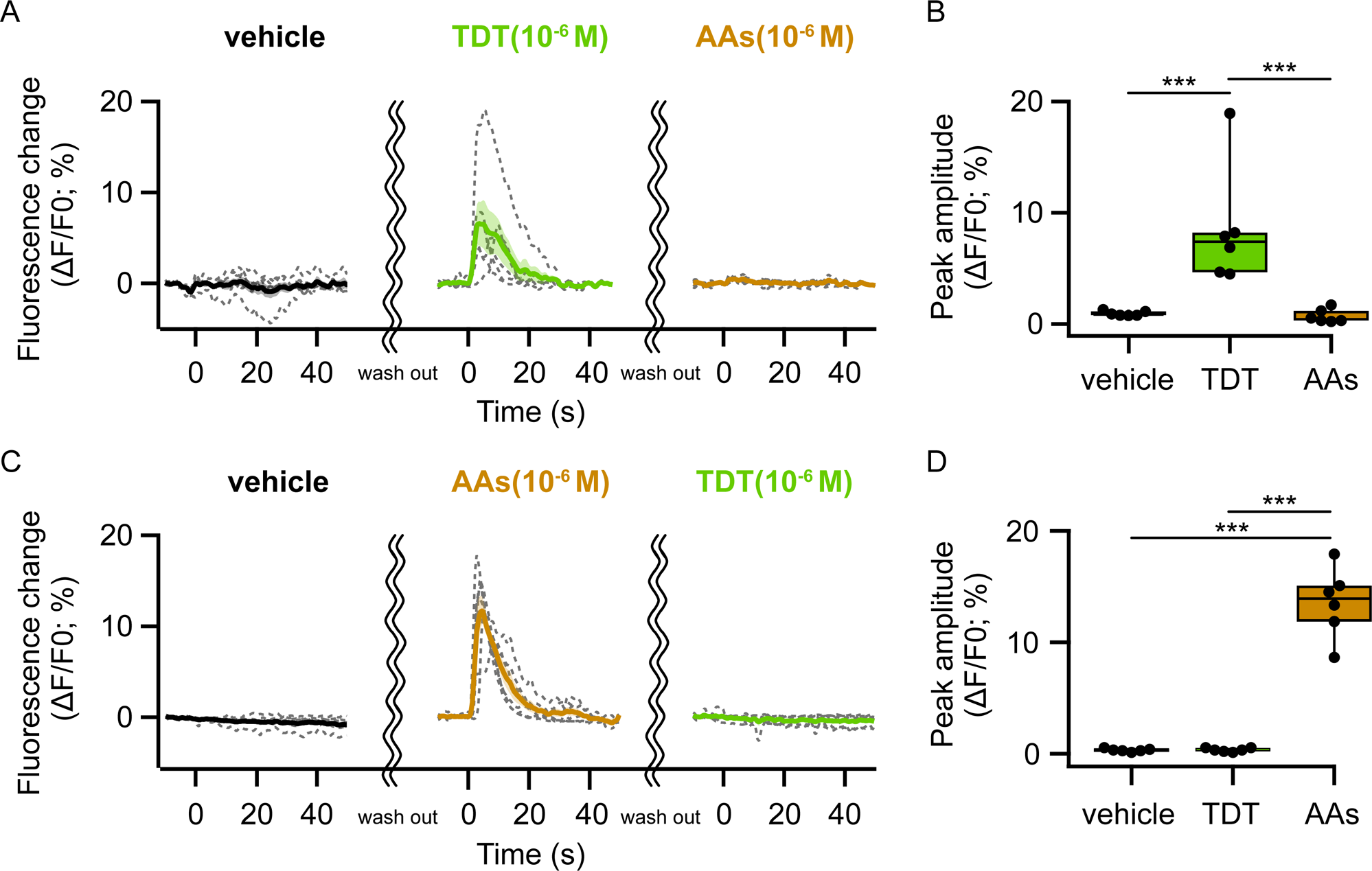
TDT-sensitive OSNs did not respond to AAs. (**A**)Fluorescence changes in TDT-sensitive OSNs to the vehicle, TDT, and AAs exposure. Individual OSNs (n = 6) are represented by gray dashed lines, with the bold line and envelope indicating the mean ± SEM. (**B**)Peak amplitude of TDT-sensitive OSNs after the administration of the vehicle (black), TDT (green), and AAs (brown). One-factor repeated measures ANOVA with Holm correction. ***P < 0.001. (**C**)Fluorescence changes in AAs-sensitive OSNs to the vehicle, AAs, and TDT. Individual OSNs (n = 6) are represented by gray dashed lines, with the bold line and envelope indicating the mean ± SEM. (**D**)Peak amplitude of AAs-sensitive OSNs after the administration of the vehicle (black), TDT (green), and AAs (brown). One-factor repeated measures ANOVA with Holm correction. ***P < 0.001.

## Discussion

In this study, we demonstrated that TDT-sensitive OSNs are present in the OE of green spotted puffers by calcium imaging of GCaMP6s fluorescence. These OSNs did not respond to AAs and TTX. The TDT-sensitive OSNs were oval-shaped cells without apical dendrites whose cell bodies were located in the superficial layer of the OE. These characteristics were similar to those of crypt OSNs previously identified by double immunohistochemistry using neuronal activity marker antibody and crypt OSN marker antibody (Suzuki et al. 2022). These results further substantiate that TDT is specifically detected by crypt OSNs.

To investigate whether live OSN of green spotted puffer detects TDT, we expressed GCaMP6s in the OSNs by *in vivo* electroporation. This technique was necessary to introduce exogenous genes to the OE of the green spotted puffer, as it is challenging to use transgenic techniques due to difficulties with lab-based breeding. The OE of green spotted puffer has notably advantageous features for *in vivo* electroporation. Generally, fish olfactory epithelium is located in nasal cavities beneath the body surface (Olivares and Schmachtenberg 2019). However, green spotted puffers have nasal organs that protrude from the dorsal head surface, exposing the OE to the external environment (cf. **Fig.6B** of Suzuki et al. 2022). This facilitates the administration of DNA solutions and the positioning of electrodes for electroporation.

The parameters of the electroporation pulse, such as waveform (exponential decay or square wave), pulse amplitude, duration, number, and frequency, are critical for optimizing gene transfer efficiency while maintaining the target cell’s physiological activity (Young and Dean 2015). We used a two-step constant-current pulse, combining a high-current, short-duration poring pulse with low-current, long-duration transfer pulses to introduce plasmid DNA into OSNs. Sadik et al. (2014) reported that a two-step pulse, combining the first large pulse and subsequent small pulses, can improve gene delivery efficiency and maintain cell viability. Another critical aspect is using constant-current pulses instead of constant-voltage pulses, which are more commonly employed in other electroporation procedures. With constant-voltage pulses, the current amplitude decays during the pulse, whereas with constant-current pulses, the driving force on the charged molecules remains constant, even if the tissue’s resistance changes during the current flow (Draghia-Akli et al. 2008). The disadvantage of electroporation is that the gene transfer efficiency is lower than other exogenous gene delivery methods. However, the sparse gene expression in the OE was advantageous for recording odorant responses from single OSNs, as it allows for better identification of individual cell morphology. For these reasons, our two-step constant-current in vivo electroporation of GCaMP6s is an adequate method for the visualization of live OSN activity.

Oval-shaped OSNs of green spotted puffers physiologically respond to TDT, not TTX. These TDT-sensitive OSNs did not respond to stimuli like AAs or the vehicle, and we found no OSNs that responded to both AAs and TDT. In teleosts, OSNs are categorized into several types, including ciliated, microvillous, crypt, kappe, and pear-shaped cells (Olivares and Schmachtenberg, 2019). This classification is based on their morphology, the position of their soma within the OE, and the specific olfactory receptor gene families they express (Kermen et al. 2013). Crypt OSNs, characterized by oval-shaped cell bodies with apical invagination, are commonly found in various fish species and positioned in the apical layer of the OE (Hansen and Finger 2000). Kappe OSNs and pear-shaped OSNs are oval OSNs reported to be located in the superficial OE only in zebrafish. Crypt OSNs can also be identified by the presence of the marker protein S100 and are notable for lack of dendrites (Hansen and Finger 2000). In contrast, kappe OSNs lack S100 but feature a characteristic cap at the apical ends of their short dendrites (Ahuja et al. 2014). Additionally, pear-shaped OSNs, characterized by their short dendrites, express the olfactory marker protein (OMP) instead of S100 (Wakisaka et al. 2017). In our study, TDT-sensitive OSNs displayed oval-shaped morphology without apical dendrite and were located on the surface of the OE. Although we did not perform immunohistochemistry for OMP, it remains plausible that the TDT-sensitive OSNs could be pear-shaped OSNs. Nonetheless, the observed morphological characteristics are well in agreement with our previous study, in which we conducted double immunohistochemistry for neuronal activity markers (phosphorated ribosomal protein S6) and crypt OSNs markers (S100) to TDT-administered OSNs (Suzuki et al. 2022). These results further substantiate that TDT is specifically detected by crypt OSNs of green spotted puffers.

Various odorants have been reported as ligands for crypt OSNs, yet the specific category of odor they respond to remains uncertain. In initial physiological experiments using acutely dispersed OE cells from the Pacific jack mackerel *Trachurus symmetricus*, AAs were reported to activate crypt OSNs (Schmachtenberg 2006; Vielma et al. 2008). A subsequent Ca^2+^-imaging study by Bazáes and Schmachtenberg (2012) reported that dispersed crypt OSNs of juvenile rainbow trout responded to a broad range of common odorants for teleosts, including AAs, bile salts, and conspecific gonadal extracts from males and females. These gonadal extracts contain reproductive pheromones. However, the crypt OSNs of sexually mature rainbow trout became responsive to only gonadal extracts from the opposite sex, and there was an increase in the number of crypt OSNs located on the OE surface. In addition, crypt OSNs respond to the kin odor, potentially linked to major histocompatibility complex II peptides, in zebrafish *Danio rerio* larva (Biechl et al. 2016), and also respond to prostaglandin E2, a pheromone that triggers synchronous spawning behavior in the surrounding individuals of adult grass puffer (Chen et al. 2022). From these results, the crypt OSNs of different fish species appear to respond to various odors, including those related to reproductive behaviors, although not exclusively. TTX (actually TDT) leaking from the oocytes has been reported to attract male grass puffers at the time of spawning (Matsumura, 1995). Therefore, the TDT is considered to satisfy the criteria for inclusion in the repertoire of biologically relevant odor ligands for crypt OSNs.

TDT is one of the major TTX analogs accumulated in toxic puffers and is considered a biosynthetic intermediate of TTX due to its structure lacking the three hydroxyl groups of TTX (Ueyama et al. 2018). Previous studies have suggested that toxic puffers prefer to feed on TTX-bearing organisms (Itoi et al. 2015, 2018, 2020; Okabe et al. 2019). These TTX-bearing organisms also contain TDT (Ito et al. 2023; Oyama et al. 2022). Furthermore, the ovaries of toxic puffers contain both TTX and TDT (Ito et al. 2023), and it is believed that they are leaking out from the cloaca (Matsumura 1995). Our previous studies (Noguchi et al. 2022; Suzuki et al. 2022) and a recent study by Miyazaki et al. (2024) demonstrated that both male and female toxic puffers are attracted to TDT. Therefore, these toxic puffers may use TDT odor as a cue to find conspecifics or to feed on TTX-bearing organisms for their efficient toxification.

## Acknowledgments

We would like to express our gratitude to Drs. Shinji Kanda (Univ. of Tokyo) and Chika Fujimori (currently, Hokkaido Univ.) for their suggestions about *in vivo* electroporation. We also thank all the staff of the Laboratory of Fish Biology for their suggestions during the experiment.

## Data availability statement

The data underlying this article will be shared on reasonable request to the corresponding author.

---

**Fig. S1:** NMR Spectra for Synthetic TDT.

**(A)** Comparison of NMR Spectra for Natural and Synthetic TDT.

**(B)** ^1^H NMR (400 MHz) Spectra of Synthetic TDT in 4% CD_3_COOD/D_2_O.

**(C)** ^13^C NMR (100 MHz) Spectra of Synthetic TDT in 4% CD_3_COOD/D_2_O.

**Fig. S2:** TDT-sensitive OSNs can be identified by their morphology before Ca^2+^ imaging.

(**A**) An depth composite epifluorescence microscopy image showing the morphology of TDT-sensitive GCaMP6s-expressed OSN before the calcium imaging experiment. Dashed white lines indicate the location of a puff micropipette. Scale bar = 5 µm.

(**B**) Fluorescence changes in TDT-sensitive OSNs to the vehicle and TDT applications. Vertical dashed lines mark the timing of odorant application.

**Fig. S3:** Repeated responses of GCaMP6s expressing OSNs to the same odorant.

(**A**) AA-sensitive OSN consistently showing Ca^2+^ responses induced by AAs.

(**B**) TDT-sensitive OSN consistently showing Ca^2+^ responses induced by TDT.

**Fig. S4:** Lack of responses in TDT-sensitive OSNs to TTX.

(**A**) Fluorescence changes in TDT-sensitive OSNs (n = 3) following administrations to the vehicle (gray), TDT (green), and TTX (purple).

(**B**) Peak amplitudes of TDT-sensitive OSNs after the administration of the vehicle (white), TDT (green), and TTX (purple). TDT-sensitive OSNs exhibit no fluorescence increase following either vehicle or TTX application (0.4 ± 0.2% for the vehicle vs. 10.0 ± 5.0% for TDT, P = 0.240; the vehicle vs. 0.4 ± 0.1% for TTX, P = 0.996; TDT vs. TTX, P = 0.240; One-factor repeated measures ANOVA with Holm correction).

