## Supplementary Information for "The odor of a nontoxic tetrodotoxin analog, 5,6,11-trideoxytetrodotoxin, is detected by specific olfactory sensory neurons of the green spotted puffers"

**A**

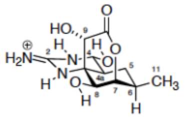

5,6,11-trideoxytetrodotoxin

| No. | synthetic [1] |  | natural [1] |  |
|---|---|---|---|---|
| | $\delta_H$ (multi., $J$ in Hz) | $\delta_C$ | $\delta_H$ (multi., $J$ in Hz) | $\delta_C$ |
| 2 |  | 155.9 |  | 155.7 |
| 4 | 5.17 (d, 9.5) | 77.5 | 5.17 (d, 10.0) | 77.3 |
| 4a | 1.99 (ddd, 13, 9.5, 3.5) | 46.3 | 1.99 (ddd, 13.3, 10.0, 3.9) | 46.2 |
| 5α | 2.08 (dt, 14.5, 4) | 27.9 | 2.07 (dt, 13.3, 4.0) | 27.8 |
| 5β | 0.92 (q, 13) |  | 0.92 (q, 13.3) |  |
| 6 | 2.13 (m) | 37.0 | 2.13 (m) | 36.9 |
| 7 | 4.61 (br s) | 87.3 | 4.61 (br t) | 87.2 |
| 8 | 4.10 (d, 2) | 74.7 | 4.10 (d, 2.3) | 74.5 |
| 8a |  | 61.3 |  | 61.2 |
| 9 | 4.63 (s) | 72.5 | 4.63 (s) | 72.4 |
| 10 |  | 177.5 |  | 177.4 |
| 11 | 1.07 (d, 6.5) | 18.4 | 1.08 (d, 6.7) | 18.3 |

All spectra were taken in 4%$CD_3COOD/D_2O$.
[1] $^1$H NMR 400 MHz and $^{13}$C NMR 100 MHz.
[2] The signal due to $CHD_2COOD$ at 2.06 ppm was used as reference for $^1$H NMR, and that due to $^{13}CD_3COOD$ at 22.4 ppm was used for $^{13}$C NMR.

**B**

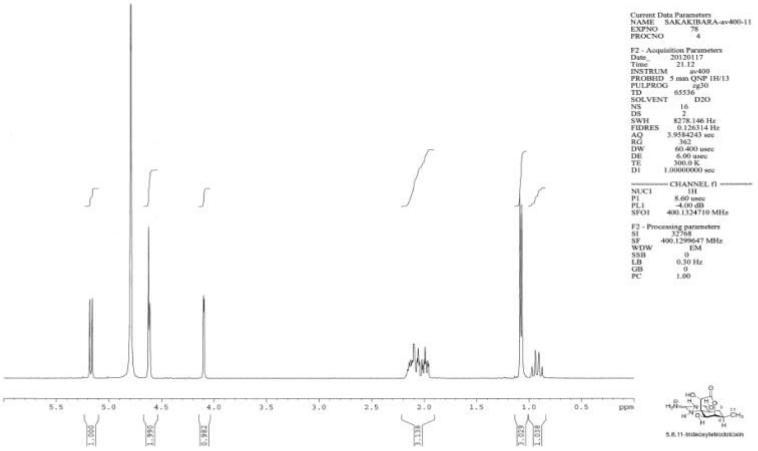

**C**

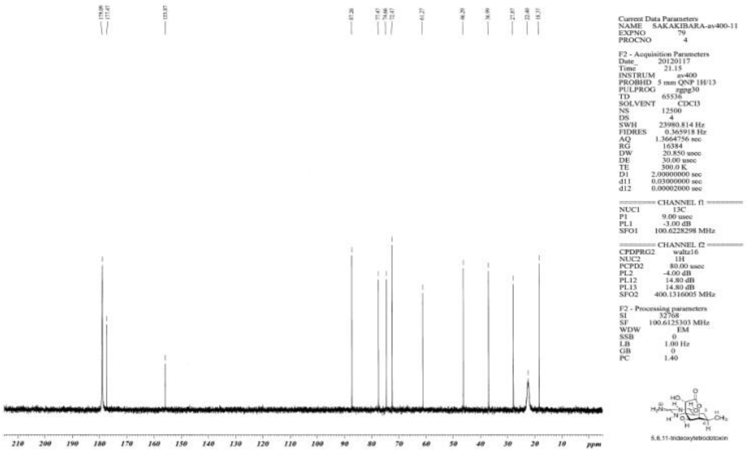

**Fig. S1:** NMR Spectra for Synthetic TDT.

**(A)** Comparison of NMR Spectra for Natural and Synthetic TDT.

**(B)** $^1$H NMR (400 MHz) Spectra of Synthetic TDT in 4% $CD_3COOD/D_2O$.

**(C)** $^{13}$C NMR (100 MHz) Spectra of Synthetic TDT in 4% $CD_3COOD/D_2O$.

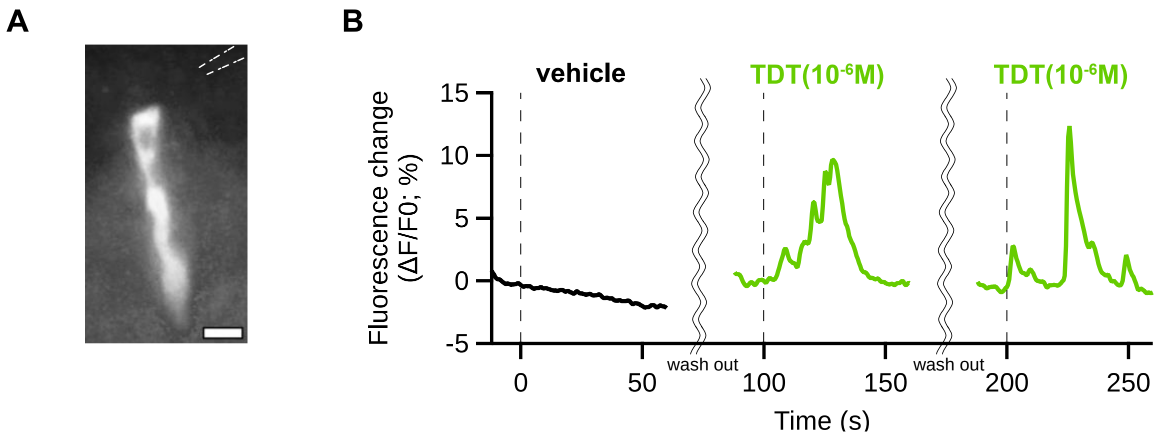


**Fig. S2:** TDT-sensitive OSNs can be identified by their morphology before $Ca^{2+}$ imaging.

(**A**) A depth composite epifluorescence microscopy image showing the morphology of TDT-sensitive GCaMP6s-expressed OSN before the calcium imaging experiment. Dashed white lines indicate the location of a puff micropipette. Scale bar = 5 µm.

(**B**) Fluorescence changes in TDT-sensitive OSNs to the vehicle and TDT applications. Vertical dashed lines mark the timing of odorant application.

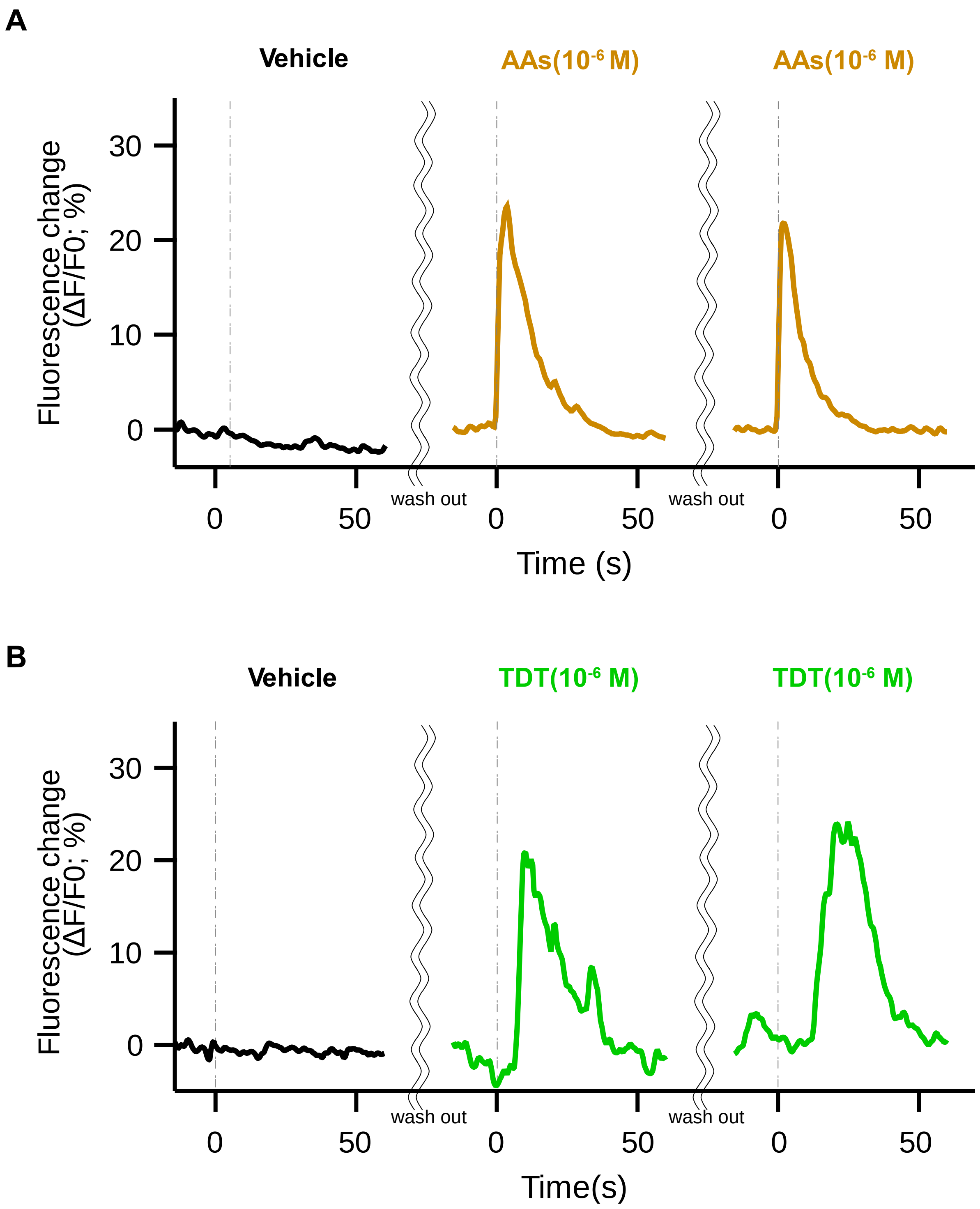


**Fig. S3:** Repeated responses of GCaMP6s expressing OSNs to the same odorant.

(**A**) An example of AA-sensitive OSN consistently showing $Ca^{2+}$ responses induced by AAs.

(**B**) An example of TDT-sensitive OSN consistently showing $Ca^{2+}$ responses induced by TDT.

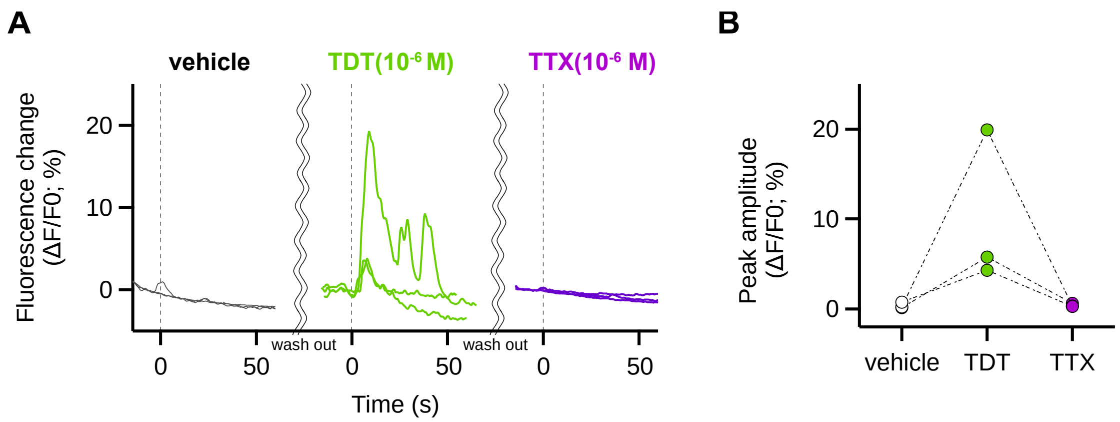


**Fig. S4**: Lack of responses in TDT-sensitive OSNs to TTX.

(**A**) Fluorescence changes in TDT-sensitive OSNs (n = 3) following administrations to the vehicle (gray), TDT (green), and TTX (purple).

(**B**) Peak amplitudes of TDT-sensitive OSNs after the administration of the vehicle (white), TDT (green), and TTX (purple). TDT-sensitive OSNs exhibit no fluorescence increase following either vehicle or TTX application (0.4 ± 0.2% for the vehicle vs. 10.0 ± 5.0% for TDT, P = 0.240; the vehicle vs. 0.4 ± 0.1% for TTX, P = 0.996; TDT vs. TTX, P = 0.240; One-factor repeated measures ANOVA with Holm correction).
